# Do examinations prepare students for higher education? A lesson from the Covid-19 lockdown

**DOI:** 10.1101/2021.01.26.428208

**Authors:** L Jones Harriet, Zini Valentina, R Green Jon, R Prendergast John, Scott Jon

## Abstract

The COVID-19 pandemic caused severe disruption to education in the UK in 2020, with most of the school teaching moving online and national school examinations being cancelled. This was particularly disruptive for those taking end of school examinations in preparation for higher education. Biological science courses require students to absorb a lot of new vocabulary and concepts, with examinations traditionally focusing on content recall rather than reasoning. Students who had entered university in September 2019 were compared with those arriving in September 2020 with respect to their knowledge of bioscience vocabulary and understanding of key concepts. Results showed no significant difference between those who had gone through the examination process in 2019 relative to those who had not, in 2020. This suggests the cramming of information for examinations has no detectable effect on the knowledge and understanding of biology that students take with them to university.

## Introduction

The transition from school or college to university is recognised as a difficult stage in a student’s education (Charalambous 2020). In 2020 the COVID-19 pandemic severely disrupted the education of students in the UK, following a rapid move to online learning and home schooling and the cancellation of A-level (and equivalent) examinations. A-level results were initially calculated using an algorithm developed by the Office of Qualifications and Examinations Regulation (Ofqual 2020a and 2020b). Following significant protest regarding the detrimental impact of this approach on the outcomes for large numbers of candidates, allocation of grades was ultimately based on Centre Assessed Grades (CAGs). The outcomes of this approach led to a significant elevation in the mean A-level grades compared with previous years (Robinson and Bunting 2020), which were then used to determine university acceptances.

The curricula for A-level (and equivalent) Biological Science courses (Department for Education 2014) include considerable content and subject-specific vocabulary. Science courses and their examinations are sometimes perceived as exercises in memory, a perception which extends into higher education (Watters and Watters 2007, Rytkönen 2012). Consequently, students are required to learn and memorise substantial amounts of material and to focus on consolidating it for examinations. Following final school/college examinations in the summer, students fail to retain much of the absorbed information by the time they arrive at university several months later (Jones et al 2015, 2019). However, students in 2020 did not go through the examination process, prompting questions about the appropriateness, efficacy and value of these methods (Cairns 2020). Students of 2020 had studied their A-level subjects for approximately 18 months before directly entering university. It has been assumed by school teachers and university lecturers that, by missing the revision phase associated with examinations, 2020 students would exhibit reduced knowledge recall compared to previous cohorts (Turner et al 2020).

The aim of the current study was to test the assumptions in the context of the 2020 student cohort’s knowledge and understanding of key biological terms. First year biology undergraduates in their first few weeks of university education were tested for biological knowledge and understanding and their results compared to a database of data from the previous year and from Jones et al (2019), when student education had not been disrupted.

## Materials and methods

An online survey comprising two sections was constructed. Section 1 focused on knowledge of vocabulary with 90 terms. This was a subset of the 476 terms previously compiled (Jones et al 2019) that were originally selected from the Letts Revise Biology (2008) study guide. The 90 terms were selected to provide a representative spread of difficulty to allow direct comparisons between two cohorts: one being pre-covid pandemic, the second in 2020 following the cancellation of end-of-school examinations. For each term in the vocabulary section, students were asked to indicate one of three choices: 1 - whether they did not know the term, 2 - whether they had heard of the term but could not explain it or 3 - whether they could explain the term. For Section 1, the 2019 cohort had 184 students, and the 2020 cohort 286 students.

Section 2 comprised 30 Multiple Choice Questions (MCQs) in which students were asked to select the correct answer from four alternatives and then state their level of confidence in their selection. The MCQs were compiled using terms from the Jones et al (2019) study and, in each question, candidates were offered a clear right answer, a vaguely right answer, a vaguely wrong answer and a very wrong answer. In the test, candidates were asked to select the clear right answer for each question then provide their level of confidence in each answer (3 = high (very confident), 2 = medium (I think this is correct), 1 = low (this is a guess). In Section 2, the 2019 cohort had 171 students, and the 2020 cohort 278 students.

First-year undergraduate students from British universities undertook the survey (anonymity meant the number of institutions is not known but exceeds four). The study was approved by the UEA ethics committee in August 2020; all answers were anonymous and student participation was voluntary.

### Statistical Analysis

To assess the effect of cohort on confidence levels an ordinal logistic mixed effects model was built using the R package “ordinal” (Christensen, 2019) which included confidence as an ordinal variable with three levels: 1, 2 or 3, as described above and included cohort as the explanatory variable. MCQ scores were calculated according to the Gardner-Medwin Certainty-Based Marking scheme (Gardner-Medwin, 2006) by combining answer correctness with confidence level to indicate levels of misconception.

Student scores were then related to study-cohort and gender using generalized linear mixed models with a Poisson error distribution. A random effect of student id and question id was used to control for pseudoreplication. For both analysis sets, the most parsimonious model was selected and considered supported if the difference in Akaike’s information criterion (AIC) value relative to the null model (ΔAIC) was greater than two units, in line with parsimony principles. All analyses were performed in R (R Core Team, 2020).

## Results

In the vocabulary test (Fig 1) and the multiple choice (MCQ) test with confidence-based marking (Fig 2), there was no significant difference in responses between the two cohorts, those pre 2020 who had sat final school examinations, and those in 2020 who had not. Looking at the mean score of correct answers in the MCQ test, out of 30 questions, the 2019 cohort scored 19.01 ± 3.67 compared to the 2020 cohort score of 19.39 ± 4.40 (error is standard deviation).

**Fig 1.**
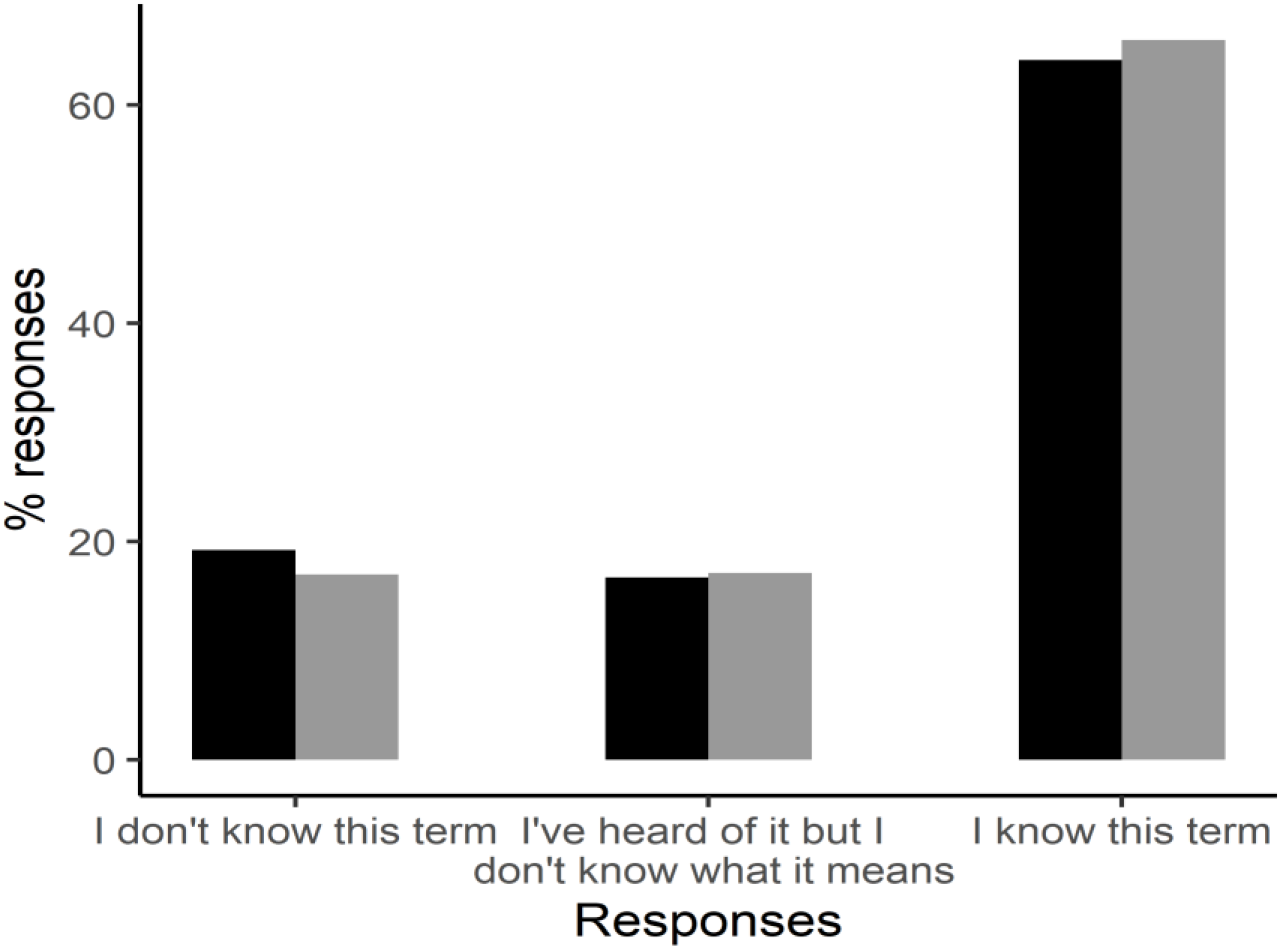
Comparison of vocabulary knowledge between the two cohorts of students. Percent responses of the 2019 cohort (black) and the 2020 cohort (grey) for the survey of their knowledge of bioscience vocabulary with three possible responses for each term.

**Fig 2.**
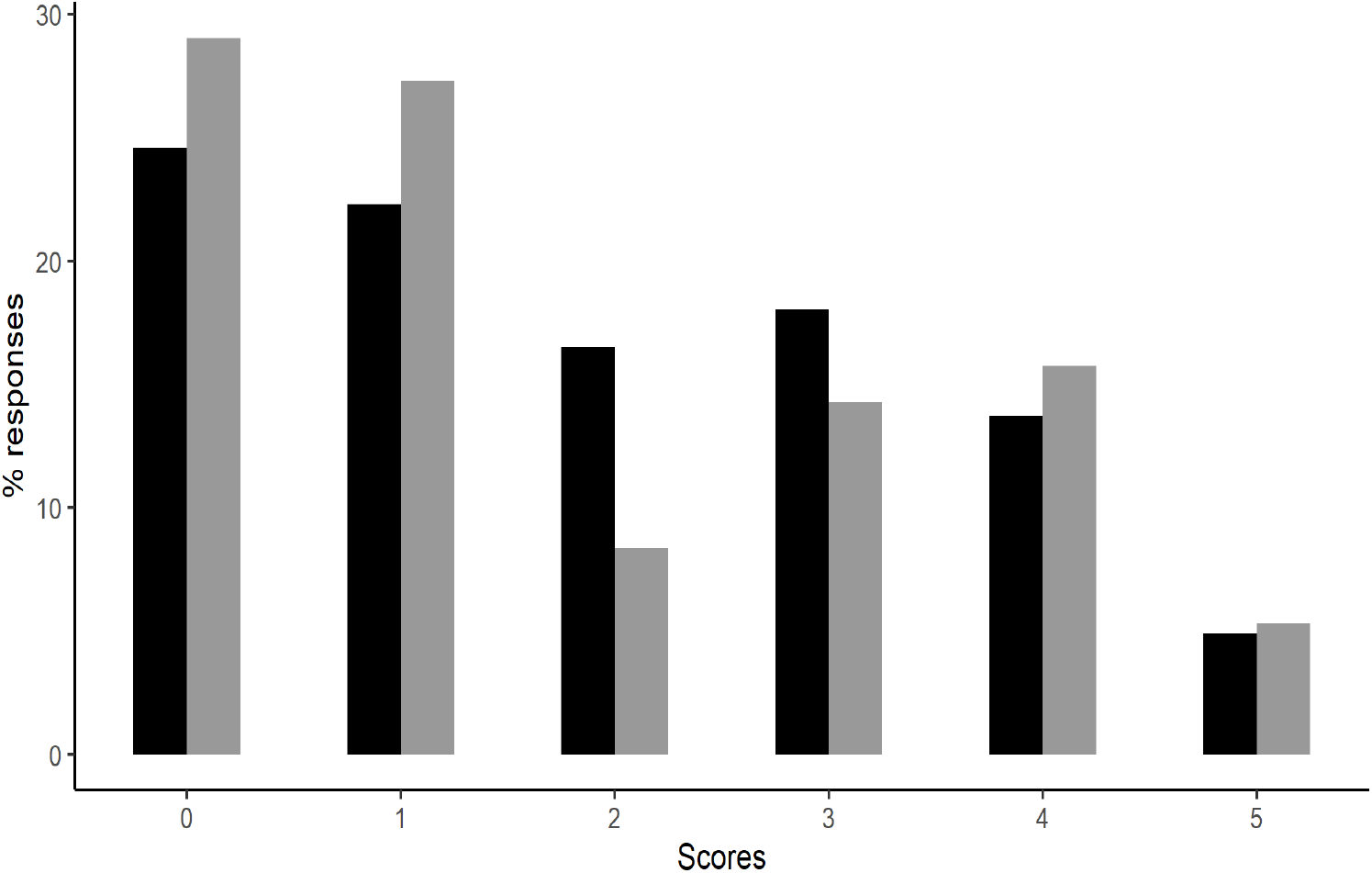
A comparison of misconceptions shown by the two cohorts of students. The percent of responses is shown from the 2019 cohort (black) and the 2020 cohort (grey) The distribution of scores ranged from 0 to 5 where 0 denotes no misconception (high confidence on the clear correct answer or medium confidence in a vaguely correct answer) and 5 denotes the most serious misconception (a high level of confidence in a very wrong answer).

Models for vocabulary and MCQ analysis show that the covid–non-covid variable was supported in neither of the models as ΔAIC with the null model was less than two units and no effect of gender on student scores was supported, where a score of greater than four would be required because of the two degrees of freedom (Table 1).

**Table 1:** Model selection of vocabulary models and MCQ. For each model AIC and the difference in AIC value relative to the best supported model (ΔAIC) are shown. Df is degrees of freedom.

| | Df | AIC | $\Delta AIC$ |
| --- | --- | --- | --- |
| **Vocabulary models** |  |  |  |
| Covid-year | 5 | 54000.63 | 0 |
| Null model | 4 | 54000.37 | 0.63 |
| **MCQ models** |  |  |  |
| Null model | 3 | 49942.39 | 0 |
| Gender | 4 | 49944.19 | 1.8 |
| Covid-year | 4 | 49944.36 | 1.97 |
| Covid+gender | 5 | 49946.17 | 3.78 |

## Discussion

In comparison to the scores of their predecessors in 2019, student knowledge of biological vocabulary and understanding of key biological concepts was found not to show any significant difference from the lack of an examination process in 2020 (Figs 1 and 2). This suggests that the examination process is not involved in the acquisition of new knowledge. Contrary to expectations (Turner et al 2020), therefore, bioscience students did not enter university this academic year (2020/21) knowing or understanding less than in previous years. Examinations are sometimes seen as a motivation for learning (Koballa and Glyn 2007) but, rather than fostering understanding, they often involve cramming many, often disjunct, items of information into short-term memory for immediate retrieval. Many of these are then forgotten by the time students reach university three months later (Jones et al 2015, 2019); although these publications showed scores for retained information did correlate with the examination grade awarded, even the top students had forgotten much of the learnt material. The current findings suggest that retained knowledge derives from the preceding 18 months of study (and possibly before) and is largely unaffected by the end-of-year grading process. The examinations are intended to provide students with an opportunity to demonstrate their knowledge, but if some of their performance is based on short term retention then it cannot be seen as assessment for learning. Assumptions are made about the value of examinations to learning (Turner et al 2020) and the relationship between the two is not simple (Lowman 1990) with over-emphasis on achievement often driving rote learning. End of school examinations should be seen only as a grading opportunity.

Students in 2020 experienced much uncertainty over the cancelling of their exams and the subsequent grading process with minority groups being particularly badly affected (Murphey and Wyness 2020). But in terms of their preparedness for university the data here show that there is little need for concern. Students, lecturers and schoolteachers need to be made aware of this to alleviate concerns of all stakeholders in the school to university transition. The 2021 and 2022 university students will, however, have missed much of the usual face-to-face, field-based and laboratory-based teaching, which data here suggest are likely to be the effective parts of their education, and these students may well need additional support. With the end-of-school grading process in doubt for 2021, focus needs to turn to student learning, as this is the best preparation for higher education, not the grading process. Tests will need to continue in future years to avoid misconceptions and potentially damaging assumptions (Turner et al 2020).

## Acknowledgements

We would like to thank Dr Ellen Bell and Dr Rebecca Lewis for their work on the MCQ test, and also Prof Stephen Rutherford, Dr Nigel Francis, Dr Sarah Ashelford, Dr Dave Lewis and any others who distributed the questionnaire to their students.

